# Sono-uncaging for Spatiotemporal Control of Chemical Reactivity

**DOI:** 10.1101/2025.06.11.659141

**Authors:** Erik Schrunk, Sunho Lee, Przemysław Dutka, Di Wu, Mikhail G. Shapiro

## Abstract

Photo-uncaging – the use of light to reveal the active part of a chemical compound by photolysis of a protecting group – has long been used to study and actuate biochemical processes. However, light scattering limits the applications of photo-uncaging in opaque specimens or tissues. Here, we introduce *sono-uncaging*, a process in which a chemical functional group becomes exposed upon the application of ultrasound, which can be applied and focused in optically opaque materials. We engineered gas vesicles (GVs), air-filled protein nanostructures sensitive to ultrasound, to contain cysteines on their concealed inner surface, hypothesizing that the application of ultrasound would collapse the GV shell and reveal the cysteines. The resulting SonoCage construct reacted with monobromobimane (mBBr), a fluorogenic, thiol-reactive molecule, only after treatment with ultrasound, establishing the sono-uncaging proof of concept. We then demonstrated the spatial patterning capability of sono-uncaging by embedding the SonoCages in an mBBr-containing hydrogel and creating fluorescent patterns with phased array ultrasound. This patterning could be accomplished using a diagnostic imaging transducer. This work establishes sono-uncaging as a method for spatiotemporal control over chemical reactivity using widely available ultrasound technology.

## INTRODUCTION

In biology, photochemical uncaging is commonly used to control the activity of ions^1–3^, neurotransmitters^4–6^, proteins and peptides^7–9^, nucleic acids^10–12^, and other bio-interactive compounds.^13,14^ However, due to light scattering and absorption, photocaged systems have limited utility in opaque biological specimens or tissues.

Unlike light, ultrasound provides centimeter-scale penetration depth into opaque materials, where it can be focused with sub-millimeter spatial precision. Ultrasound has been used to control chemical reactions via cavitation-driven mechanical forces on mechanophores^15–17^ or the generation of reactive oxygen species.^17–20^ However, the conditions commonly used to elicit cavitation effects can be harmful to tissues and require specialized focused ultrasound equipment^15,21^, limiting the utility of existing ultrasound-triggered chemistries in living systems.

In this study, we set out to establish control of biochemical functions using mild, non-cavitating ultrasound that can be applied using common imaging devices (**Figure 1a**). Our approach takes advantage of gas vesicles (GVs), genetically encodable air-filled protein nanostructures (∼85 nm diameter, ∼500 nm length)^22^ that have recently emerged as the first biomolecular agents for ultrasound imaging and actuation (**Figure 1b**). The stark density and compressibility differences between GVs’ air-filled interiors and their aqueous surroundings endow these nanoparticles with strong ultrasound contrast.

**Figure 1.**
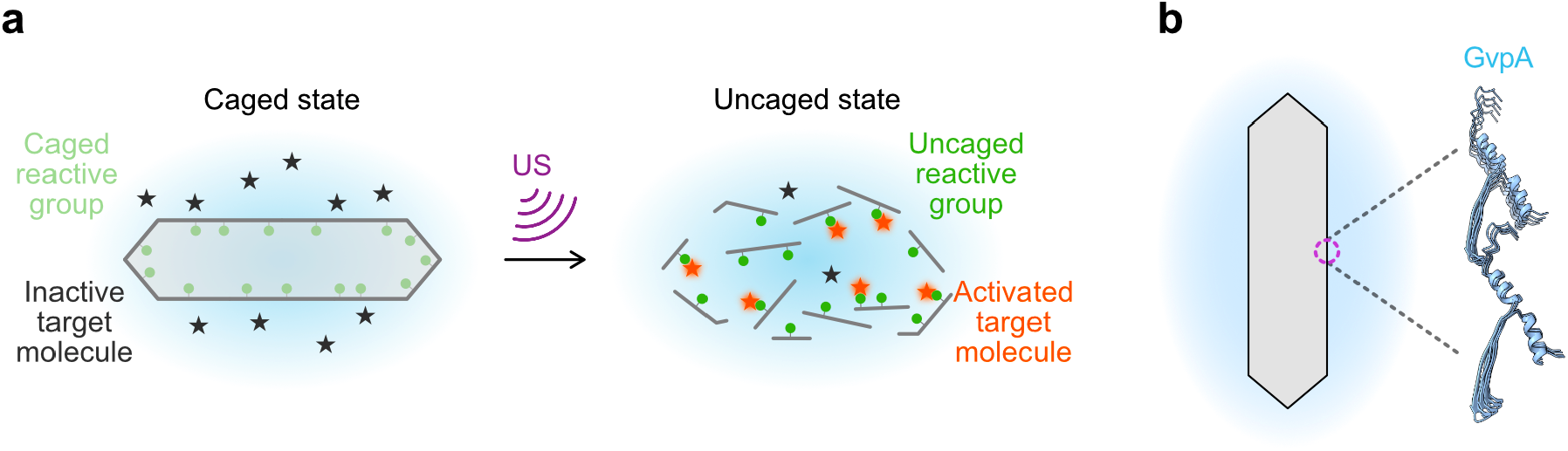
Sono-uncaging and SonoCages. (a) Schematic of sono-uncaging. The caged state encloses the reactive group (green dot) within the air-filled interior of the SonoCage. A target molecule (star) that reacts with the caged group is in solution, physically separated from the caged groups by the GV walls. After application of ultrasound (US), the reactive group is uncaged and free to react with the target molecule. (Reacted or “activated” target molecule depicted in orange.) (b) SonoCages are engineered from GVs, which are composed of repeating units of the structural protein GvpA.

Meanwhile, their biocompatibility, engineerability and genetic encodability have enabled their use in a variety of biological scenarios.^23–30^ One aspect of GVs’ interaction with ultrasound is that they mechanically collapse under acoustic pressure above a certain pressure threshold.^23,28,31^ This collapse results in the partial exposure of their shell interior, which is normally shielded from aqueous media.

Here, we describe SonoCages, which are GVs engineered to have a concealed cysteine group facing the interior of the GV that, upon a mild ultrasound exposure, reveal a reactive reduced thiol functional group—a process we call “sono-uncaging.” The GV shell is made primarily of repeating subunits of the structural protein GvpA (**Figure 1b**), which we engineered via a combination of structural predictions and a site-scanning mutagenesis library to introduce a well-tolerated interior-facing cysteine substitution. The resulting SonoCages can undergo ultrasound-activated exposure of cysteine and turn on monobromobimane^32,33^ (mBBr) fluorescence, thereby demonstrating the sono-uncaging proof of concept. Furthermore, using a phased-array imaging transducer, we are able to pattern chemical reactivity in hydrogels containing SonoCages, resulting in the turn-on of mBBr fluorescence in several digitally programmed patterns. Thus, these SonoCages represent a spatially, temporally controllable source of chemical reactivity that can be activated by ultrasound.

## RESULTS AND DISCUSSION

### Several internal-facing residues in GvpA1 are tolerant to mutations to cysteine

To develop sono-uncageable GVs, we sought to introduce a unique chemical group into GVs’ interior walls such that the chemical moiety would only be exposed once the GVs are collapsed. We chose to uncage cysteine for several reasons. First, as a natural amino acid, cysteine can be readily incorporated into the GV shell by endogenous translation machinery. Second, of the natural amino acids, cysteine is the only molecule to possess the thiol functional group, which participates in rather unique chemical reactions (thiol-maleimide, thiol-iodoacetamide, etc.) compared to other amino acids, allowing for cysteine-specific applications and assays. Finally, wild-type (WT) GvpA does not contain cysteine, meaning that WT GVs are not able to undergo sono-uncaging of cysteine and can serve as cysteine-free controls.

To introduce cysteine into the GVs’ interior walls, we cross-referenced predicted interior-facing residues of GvpA with experimentally determined cysteine-tolerant amino acid sites to arrive at a short list of potential sono-uncaging candidates. Working with the bARG_Ser_ GV gene cluster_34_, we synthesized and cloned a cysteine scanning mutant library of the *gvpA1* gene and transformed it in bulk into *E. coli*, which we then plated onto inducer-containing agar dishes. We used the opacity of induced colonies as a proxy for GV expression (as GV-expressing colonies look white) to identify and sequence the cysteine mutations that do not abrogate GV expression. Then we compared these cysteine-tolerant sites with the positions of side chains predicted to be inward-facing (**Figure 2a**) from the structural model of GvpA^22,35^ (from *Anabaena flos-aquae*, whose GvpA has 92% similarity to our GvpA1^26^). This comparison yielded four mutants—V16C, V17C, V33C, and V47C—that both expressed GVs and were expected to have interior-facing thiol groups (**Figure 2b-d**).

**Figure 2.**
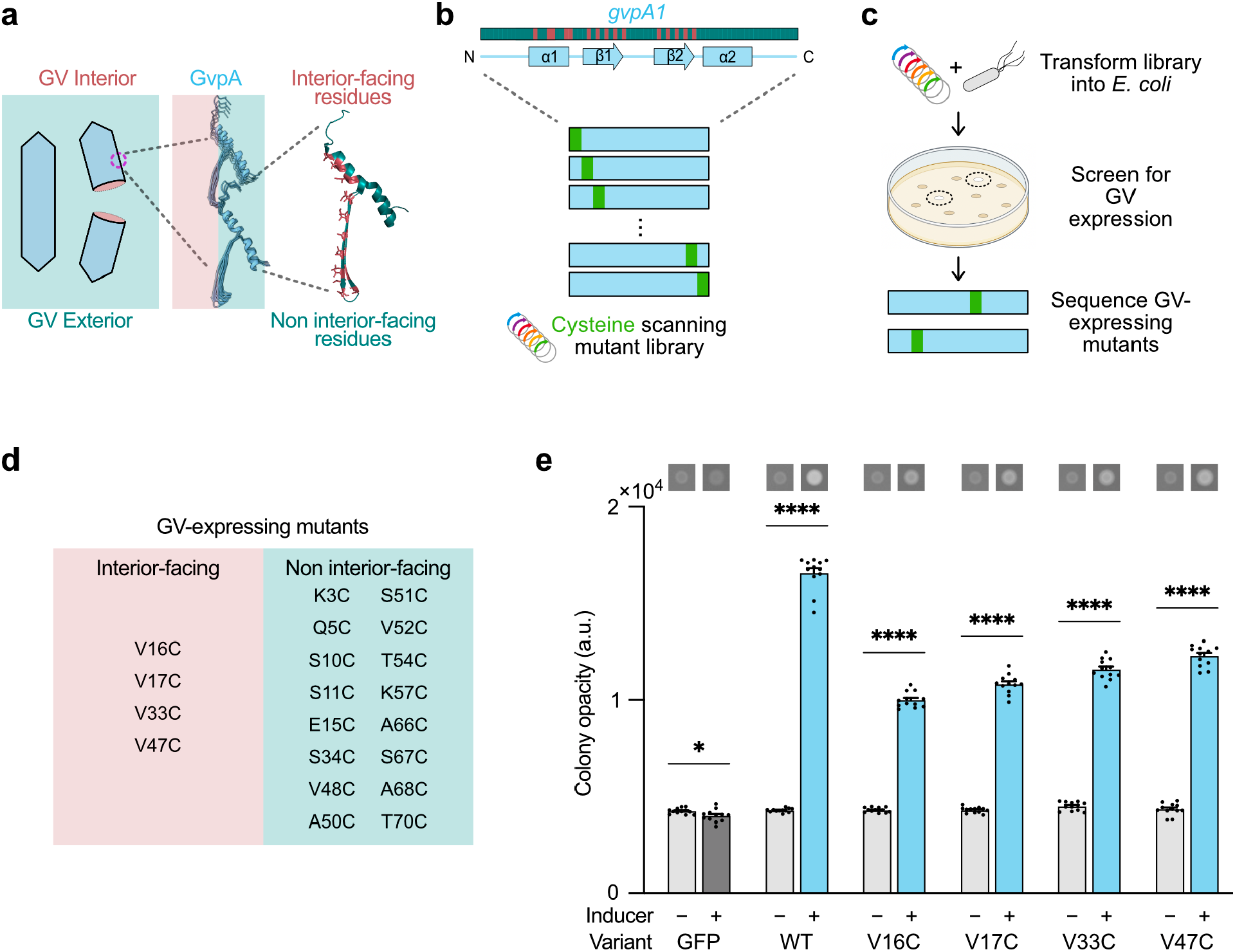
GvpA cysteine scanning library identifies several SonoCage candidates with potentially inward-facing cysteine side chains. (a) The GvpA that makes up GV shells borders both the air-filled GV interior (pink) and the aqueous GV exterior (teal). In the diagram of GvpA at the right, residues of GvpA with predicted interior-facing side chains are shaded in pink, and the rest in teal. (b) A cysteine scanning mutant library for the *gvpA1* gene that codes for GvpA1, the primary structural protein in GVs. Linear diagram showing the relative positions of the interior-facing (pink) and non interior-facing residues (teal) shown above. (c) Workflow for the development of a cysteine-enclosing SonoCage. The cysteine scanning mutant library was transformed into *E. coli* and GV expression was induced. Colonies that turned white were treated as GV-expressing and sequenced. (d) Diagram depicting the predicted side chain orientation of all GV-expressing cysteine mutants discovered. Candidates for sono-uncaging of cysteine are on the left. (e) Graphs of opacity (a proxy for GV expression) in bacterial patches transformed with plasmids encoding sono-uncaging candidate and WT GV expression. A plasmid encoding GFP expression is also included as a GV-negative control. Induced and uninduced patches are grouped; a representative image of each condition is shown above its corresponding column. N = 12 patches per condition. Patches with a plasmid encoding green fluorescent protein (GFP) expression were included as a GV-negative control. Asterisks represent statistical significance by unpaired t-tests (****: p<0.0001, *: p<0.05). Error bars represent mean ± SEM. The V33C mutant is a double mutant, with an additional mutation at A50.

To compare their relative expression levels, we expressed all four mutant GV variants alongside WT GVs and green fluorescent protein (GFP) in replicate bacterial patches and measured their optical opacity (**Figure 2e, S1**). We moved forward with the best-expressing variant, V47C, as our top candidate for SonoCages.

### V47C mutant GVs demonstrate sono-uncaging in response to collapse triggered by ultrasound

After selecting a putative SonoCage variant, we tested its capacity for the uncaging of cysteine reactivity upon exposure to ultrasound.

We incubated purified V47C mutant GVs with monobromobimane (mBBr), a molecule that reacts with reduced thiol under biological conditions and drastically increases fluorescence, and applied ultrasound using a Richmar Soundcare Plus device, which operates at 3 MHz and 2 W/cm^2^ intensity. We observed that the V47C GVs produced a very large relative increase in mBBr fluorescence after the application of ultrasound compared to WT GV controls in phosphate-buffered saline media (**Figure 3b**). The slight increase between ultrasound-treated and non-treated WT GV samples can be attributed to a drop in opacity (and increase in brightness) concomitant with GV collapse. Confocal microscopy similarly revealed that only the ultrasound-treated V47C sample exhibited higher-than-background mBBr fluorescence (**Figure 3d**). These results confirmed that our mutant GVs could undergo sono-uncaging—we had indeed developed SonoCages. In addition to working in solution, we found that sono-uncaging was effective when the GVs were embedded in hydrogel media (0.5% low-melt agarose) (**Figure 3c**).

**Figure 3.**
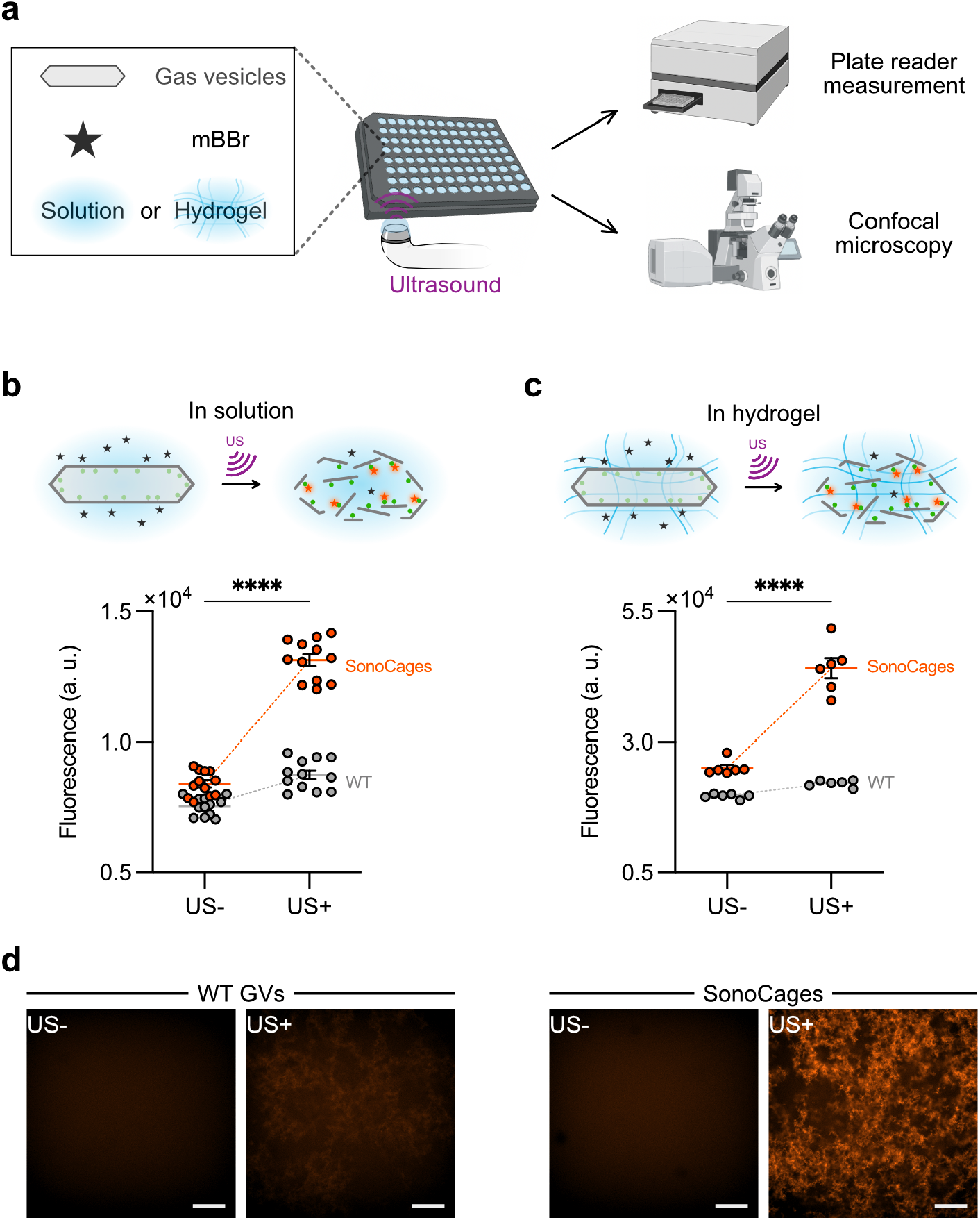
SonoCages (V47C mutant GVs) undergo sono-uncaging of cysteine in solution and in agarose hydrogels. (a) Schematic of process flow for sono-uncaging proof-of-concept experiments. mBBr fluorescence was assessed with a plate reader (b and c) and confocal microscope (d). (b) and (c) Graphs of mBBr fluorescence for WT GVs and SonoCages in PBS (b) or 0.5% w/v low-melt agarose (c) with and without ultrasound (US). N = 12 (b) and N = 6 (c) wells of a 96-well plate per condition. Asterisks represent statistical significance by difference-in-differences (****: p<0.0001). Mean values are depicted as horizontal bars. Error bars represent mean ± SEM. (d) Confocal fluorescence microscopy images of WT GVs and SonoCages in PBS. Representative z-slices shown. All scale bars 200 µm. Fluorescence is highest in the ultrasound-treated SonoCage samples due to the mBBr-thiol reaction.

### SonoCages allow for spatiotemporal control of thiol reactivity

After validating the sono-uncaging proof of concept with our SonoCages, we demonstrated their ability to work in conjunction with ultrasound to create desired spatial patterns of chemical reactivity in 3 dimensions. We prepared a 0.25% agarose hydrogel containing SonoCages and used ultrasound to collapse the GVs in a pattern (**Figure 4a**).

**Figure 4.**
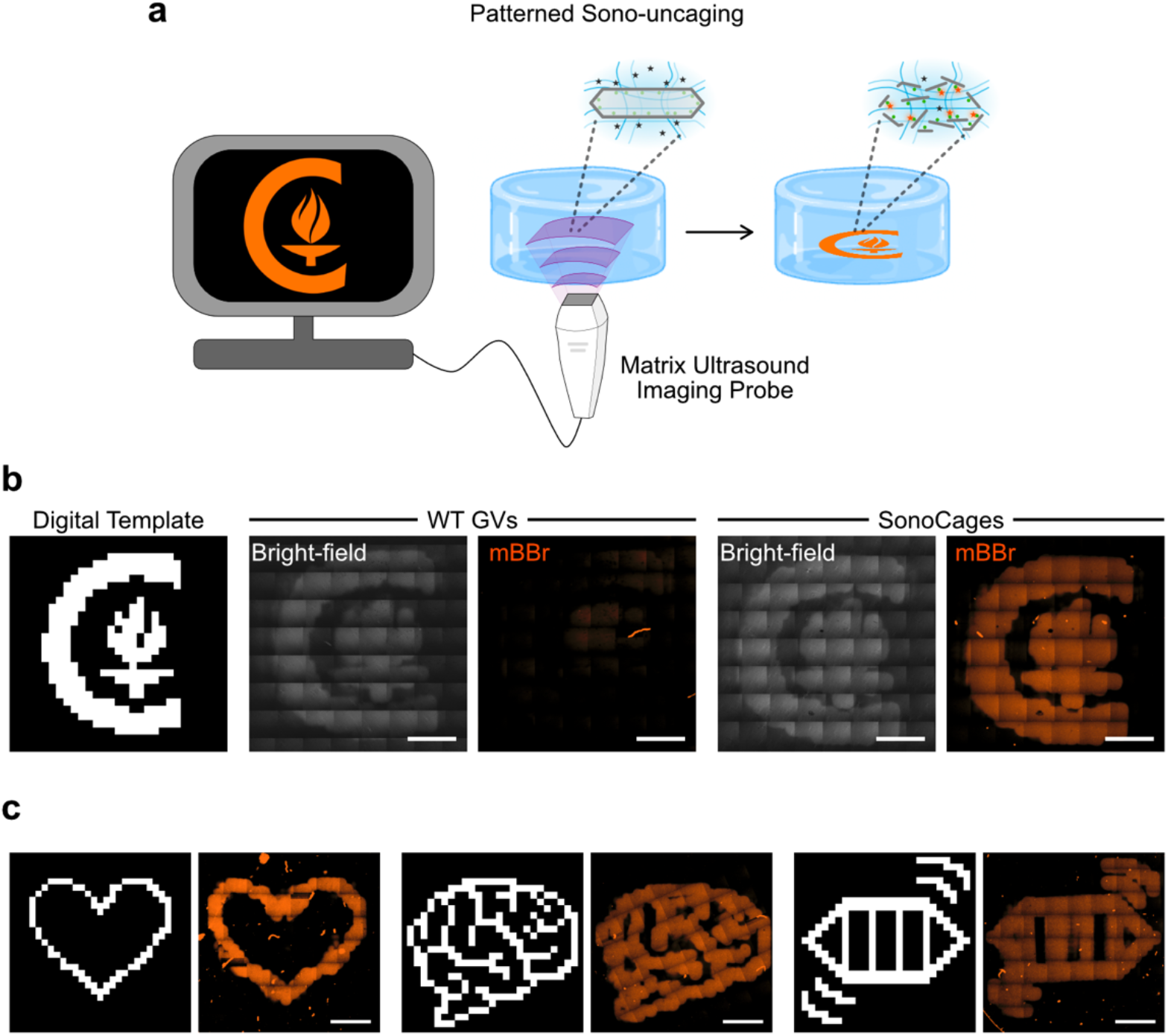
Sono-uncaging of SonoCages leads to patterning of thiol reactivity with spatiotemporal control in an agarose gel. (a) Schematic for ultrasound patterning of thiol reactivity. An agarose hydrogel containing intact SonoCages and mBBr was prepared. Ultrasound was used to create a Caltech logo pattern of collapsed GVs, leading to thiol exposure and reaction with mBBr only in areas of GV collapse. (b) Confocal images of Caltech logo patterns in 0.25% low-melt agarose with 10 µM mBBr and either WT GVs or SonoCages. All scale bars 2 mm. (c) Confocal mBBr fluorescence images of SonoCage samples with other patterns. All scale bars 2 mm.

To create our patterns, we generated a focused beam (transmit frequency = 8.9 MHz, focal length = 7 mm, angle = 0°) using 8 × 8 elements from a diagnostic matrix array ultrasound transducer (32 × 32 elements, pitch size = 0.3 mm) which we electronically steered across the samples based on patterns drawn on a 25 × 25 binary grid. The peak positive and negative ultrasound pressures at the focal point were 2.2 and −1.5 MPa, respectively. At each location, the ultrasound was applied for a total of just ∼1 µs (number of cycles = 5).

Visualizing the resulting activated hydrogels by confocal microscopy, we found mBBr fluorescence patterns that closely corresponded to our desired shapes (**Figure 4b**). While regions of patterned GV collapse could be observed in both SonoCage and WT GV samples, the WT control did not exhibit patterned fluorescence, confirming the specificity of our uncaging reaction. We implemented several other patterns, all of which behaved similarly (**Figure 4c**). These results underscore the advantage of sono-uncaging: mild ultrasound effected a covalent chemical reaction in a predetermined digitally programmed pattern. While in this experiment fluorescence was used as the readout for the thiol-mBBr reaction, this same approach could be used to pattern thiol reactivity in opaque media and living tissues.

## CONCLUSION

Taken together, these results illustrate the use of mild, bio-compatible ultrasound to exert spatiotemporal control over chemical reactivity with SonoCages. Our cysteine mutant library screen revealed that at least 27% of amino acids in GvpA could individually be mutated to cysteine without abrogating GV expression, with four of these sites corresponding to likely inward facing side chains. The best-expressing V47C variant produced GV SonoCages that reacted with a fluorogenic, thiol-reactive compound specifically after application of ultrasound. Furthermore, we used mild ultrasound from a diagnostic matrix transducer to create millimeter-scale patterns in a SonoCage-containing hydrogel, and found that fluorescence, and thus thiol reactivity, was localized to the intended locations. While in this study we demonstrated the sono-uncaging of thiol, we anticipate that this approach can be generalized to other reactive amino acid side chains. Given the vast range of chemical groups in non-canonical amino acids^36^, sono-uncaging could be used to trigger a variety of reactions for diverse applications; a simple mutant expression screen in an appropriate bacterial strain could yield the desired SonoCage.

As they are based on GVs, SonoCages could be imaged with ultrasound or other modalities prior to their collapse, and these imaging methods could also confirm that collapse, and therefore uncaging, has taken place. Furthermore, GV engineering could endow sonouncaging with additional capabilities, such as serial actuation of multiple functionalities with different acoustic pressures.^37^ With such future extensions, SonoCages establish a versatile platform for controlling chemical reactivity with mild ultrasound.

## EXPERIMENTAL METHODS

### Design, assembly, and screening of GvpA1 cysteine scanning mutant library

The cysteine scanning GvpA1 library was constructed in a manner similar to the scanning site saturation libraries described by Hurt et al.^27^ except that each amino acid in GvpA1 was mutated to only cysteine. The arabinose-inducible bARG_Ser_ plasmid_34_, which contains the *gvpA1* gene under the control of the pBAD promoter, was used as the starting point for the mutant library construction. *gvpA1* was divided into three subunits that tiled the gene, and three oligo libraries were designed such that each library contained a middle variable region covering one subunit of the gene plus two invariable anchoring regions on either side of the variable region. The variable region substituted each codon individually to TGT (cysteine); the anchoring regions were used as primer binding sites in the PCR amplification of the oligo libraries and as overlapping regions in the Gibson assembly of the mutant plasmids as described below. These oligo libraries were synthesized as one oligo pool by Twist Bioscience (South San Francisco, CA). The three oligo libraries were then amplified from the pool with PCR using primers designed for each library’s anchoring regions. Only 14 total cycles of PCR were used to lower the risks of mutation swapping and of any one oligo dominating the library. The amplified oligo libraries were isolated by electrophoresis using a 3% agarose gel with TBE (tris-borate-EDTA) buffer and purified with a New England Biolabs (“NEB;” Ipswich, MA) DNA cleanup column. The three oligo pools were assembled into mutant plasmid libraries via Gibson assembly (NEB); the anchoring regions of the oligos served as the overlaps between the oligos and the corresponding backbone fragments as well as the binding sites of the PCR primers used to amplify those backbone fragments. The three mutant plasmid libraries were then transformed into NEB Stable electrically competent cells and plated onto solid agar media in either inducing (1% w/v L-arabinose and 0.1% w/v glucose) or non-inducing (1% glucose, no arabinose) conditions. Colonies that appeared white (indicating GV expression) on the induced plates were miniprepped and sequenced.

### GV interior-cysteine candidate expression screen

GV-expressing cysteine variants identified in the cysteine scanning library were then cross-referenced with GvpA1 amino acid sites believed to be interior-facing (**Figure 2d**), leading to four putative interior-cysteine candidates. The candidate variants’ relative GV expression levels were compared using an opacity-based assay of GV-expressing bacterial patches as described in our previous work.^26^ Briefly, *E. coli* transformed with the four mutant sono-uncaging candidate plasmids, as well as control plasmids encoding WT GvpA1 and GFP under the same promoter, were plated in patch format by resuspending a colony in 100 µL of phosphate-buffered saline (PBS) and depositing 1 µL of that suspension onto inducer LB media plates containing 1.5% (w/v) agar, 1% (w/v) L-arabinose, 0.1% (w/v) glucose, and 25 µg/mL chloramphenicol as well as arabinose-free control plates with 1% (w/v) glucose, agar, and chloramphenicol. For each plasmid, a total of twelve induced and twelve uninduced patches were made—two distinct colonies were each used to make three replicate patches on two induced and two uninduced plates—and grown at 37°C for 1 day. GV expression was quantified using a ChemiDoc gel imager (Bio-Rad; Hercules, CA) by measuring the opacity of the patches as a proxy for expression (Figure S1). Images were processed using ImageJ (NIH; Bethesda, MD). The screen revealed that the V47C mutant expressed the best of the variants (**Figure 2e**).

### Large-scale expression and purification of V47C and WT GVs

GVs were expressed and purified using a protocol similar to that described by Lakshmanan et al.^38^ for the heterologous expression and processing of *Bacillus Megaterium* GVs in *E. coli*.

BL21(DE3) electrically competent *E. coli* cells (NEB) were transformed with the bARG_Ser_ plasmid with either WT or V47C mutant *gvpA1* and plated. Individual colonies of each variant (WT or V47C) were picked and grown overnight in LB media with 25 μg/mL chloramphenicol and 1% w/v glucose. The next day, the cultures were saved as glycerol stocks; those glycerol stocks were used to seed all GV-expressing cultures.

GVs were expressed in large batches using 250-mL baffled flasks containing 35 mL autoinduction medium (LB liquid medium with 0.3% w/v glucose, 0.5% w/v L-arabinose, 0.6% w/v glycerol, and 25 μg/mL chloramphenicol). A 200 μL pipette tip was dug into the appropriate glycerol stock to excavate a small amount (∼10 μL) of the frozen stock; this frozen stock was then added into 1.5 mL of media in a tube and pipetted up and down. 200 μL of this mixture was used to seed each of six flasks. Once cells were added to each flask, the flasks were grown at 37°C with 250 rpm shaking for 24 hours.

GV-expressing cultures were poured into 50-mL centrifuge tubes (35 mL of liquid per tube) and centrifuged overnight at 4°C and 350g. The next day, the GV-expressing cells had formed a buoyant layer on top of each tube. The pellet and as much of the media as possible were removed from each tube using an 18G needle and syringe, and then the cells were lysed by resuspending in Solulyse bacterial protein extraction reagent (AMSBIO; Cambridge, MA) totaling 50 mL per variant with 250 μg/mL lysozyme and 10 μg/mL DNase I added. The tubes were rotated at 37°C for 2 hours at 10 rpm, then centrifuged at 4°C and 350g for 8 hours in 50-mL tubes not exceeding 35 mL liquid volume each. This lysis step was performed a total of four times, but with 25 mL total Solulyse (with lysozyme and DNase I) per variant instead of 50 mL for the subsequent lysis steps. Next, the lysed GV suspension was washed by removing the liquid and pellet from each centrifuged tube and resuspending in 25 mL PBS, then centrifuging the tubes at 4°C and 350g for ∼8 hours. This PBS washing step was performed a total of four times as well. After the fourth PBS wash, GVs were unclustered by resuspending them in ∼8 mL of 6 M urea in PBS, splitting them into 2-mL tubes with 1 mL volume each, and rotating these tubes at room temperature and 10 rpm for 2 hours. The GVs were then centrifuged for ∼6 hours at 4°C and 250g. This unclustering step (removal of liquid with needle, resuspension in 6 M urea in PBS, incubation with rotation, and centrifugation) was also performed a total of four times. Finally, the GVs were dialyzed against 4 L PBS in 6-8 kDa MWCO dialysis tubing a total of four times (i.e. with three buffer changes) to rid them of urea. This resulted in roughly 10 mL of concentrated GVs (V47C OD_500_ of ∼10, WT OD_500_ of ∼18, where “OD_500_” refers to the optical depth of the suspension at 500 nm). Throughout this process, care was taken not to shake or drop tubes containing GVs to protect them from collapse. Vigorous pipetting of GVs during resuspension steps appears to be tolerated.

### Sono-uncaging proof-of-concept experiment

A 96-well plate was loaded with GVs (V47C or WT) or PBS (as a control) mixed with monobromobimane (mBBr) (**Figure 3b**) and additional low-melt agarose (LMA, **Figure 3c**). The final concentrations were OD_500_ 3 GVs, 10 μM mBBr, and 0.5% LMA (when used) in 100 μL total volume per well. Each well was loaded by mixing concentrated GVs (or PBS) with concentrated mBBr or a concentrated mBBr-LMA mixture as described below.

Purified GV stocks (V47C and WT) were diluted 100x in PBS (10 μL GV stock into 990 μL PBS) and their OD_500_ values measured using a cuvette with a 1 cm path length in a Thermo Scientific NanoDrop 2000 spectrophotometer (Thermo Scientific; Waltham, MA). The approximate stock OD_500_ values were then back-estimated from the diluted OD_500_ values by multiplying by 100. Wells were loaded with concentrated GVs such that the final GV OD_500_ would be 3.

In the experiments without LMA, concentrated mBBr was added into the wells immediately before administration of ultrasound (see below) for a final concentration of 10 µM; the wells were then pipetted up and down to mix. mBBr was purchased from Millipore Sigma (Burlington, MA) as a powder (THIOLYTE, 596105) and reconstituted at a concentration of 13 mg/mL (50 mM) in acetonitrile weeks prior to the experiment; it was aliquoted and stored at 4°C until used.

In the experiments with LMA, a concentrated LMA-mBBr mixture was prepared to reduce the number of pipetting steps. LMA was prepared at a 2% w/v concentration in PBS several days before the experiment by adding 2 g of LMA (GoldBio; St. Louis, MO) into 100 mL PBS and microwaving until the agarose dissolved. The LMA solution was stored at 70°C for several days to degas. On the day of the experiment, a 1.25% LMA solution containing 25 μM mBBr was prepared by adding 750 μL PBS to a 2-mL tube and then pouring 2% LMA solution up to the 2-mL line as 2% LMA is difficult to pipette. The tube was then inverted several times to mix and stored in a heat block at 42°C. 1 μL of a 50 mM mBBr stock solution was added to the tube and the tube inverted to mix, resulting in a final concentration of 1.25% LMA and 25 μM mBBr. Each well of the plate was loaded with 60 μL OD_500_ 5 GVs and 40 µL of this LMA-mBBr solution and pipetted up and down 5 times to mix, resulting in the desired final concentrations.

Immediately after each well was loaded, half of the wells were placed onto a handheld therapeutic transducer (Richmar Soundcare Plus; Clayton, MO) operating at 3 MHz and coupled with ultrasound gel. The plate was moved over the transducer, resulting in the collapse of the GVs in half of the wells, which visibly lost their opacity compared to the un-collapsed wells. Following ultrasound collapse, the plate was wiped with a tissue to remove residual ultrasound gel and stored in the dark for 1 hour. After 1 hour, the wells of the plate were measured with a Tecan Spark (Tecan Group; Zürich, Switzerland) fluorescence reader with 380/480 nm excitation and emission.

### Ultrasound patterning of GV collapse

For the ultrasound patterning experiments, samples were prepared on individual mylar-bottomed culture dishes. The dishes were made prior to the experiment by removing the glass bottoms from 35 mm dishes (Matsunami Glass USA; Bellingham, WA) and gluing mylar films to the bottoms of the dishes.

On the day of the experiment, the dishes were loaded with GVs, LMA, and mBBr totaling 200 µL, just enough volume to evenly coat the entire inset well of each dish. The dishes were prepared to a final concentration of OD_500_ 3 GVs (WT or SonoCages), 0.25% LMA, and 10 µM mBBr, but were loaded in a slightly different manner as before due to the logistical differences between setting up a multiwell assay and individual dishes. In brief, each plate was first loaded with concentrated GVs (WT or SonoCages), then diluted with PBS to 98 µL. Next, 2 µL of a 1 mM mBBr solution was added. Finally, 100 µL of a 0.5% LMA solution in PBS (prepared by pouring 500 µL 2% LMA stock solution into 1500 µL PBS in a 2-mL tube and storing at 42°C until ready for use) was added; the mixture was then pipetted up and down, which served both to mix the components and evenly spread them on the mylar. Once the samples were prepared, they were left in the dark for around 15 minutes to solidify, then patterned with ultrasound in a water tank.

After the plate was placed on the water surface, a focused ultrasound beam was transmitted onto the plate following pre-designed patterns using a 1024-element matrix array probe (Vermon; Tours, France; center frequency = 15 MHz, pitch size = 0.3 mm). The probe was submerged in water degassed with a water conditioner (ONDA; Sunnyvale, CA) and positioned at a 7 mm focal distance from the bottom of the plate. The probe’s distance to the plate and its horizontality were confirmed through B-mode imaging. An axially directed focused beam was generated using an 8 × 8 subset of transducer elements from the full 32 × 32 array, enabling the printing of arbitrary patterns based on 25 × 25 two-dimensional binary masks. The transmit frequency, duty cycle, and the number of cycles were set to 8.9 MHz, 0.67, and 5, respectively. The pulse repetition frequency was 4 kHz. The input voltages applied to the transducer were 25 V for WT GV samples and 20 V for SonoCage samples.

After ultrasound patterning, the dishes were imaged with a Zeiss LSM 800 confocal microscope with ZEN Blue. Images were processed with the Fiji package of ImageJ.

## ACKNOWLEDGEMENTS

The authors would like to thank Dr. Andres Collazo and Dr. Giada Spigolon of the Biological Imaging Facility (BIF). Imaging was performed at the BIF with the support of the Caltech Beckman Institute and the Arnold and Mabel Beckman Foundation. M.G.S. is an investigator of the Howard Hughes Medical Institute.

## SUPPORTING INFORMATION

**Figure S1.**
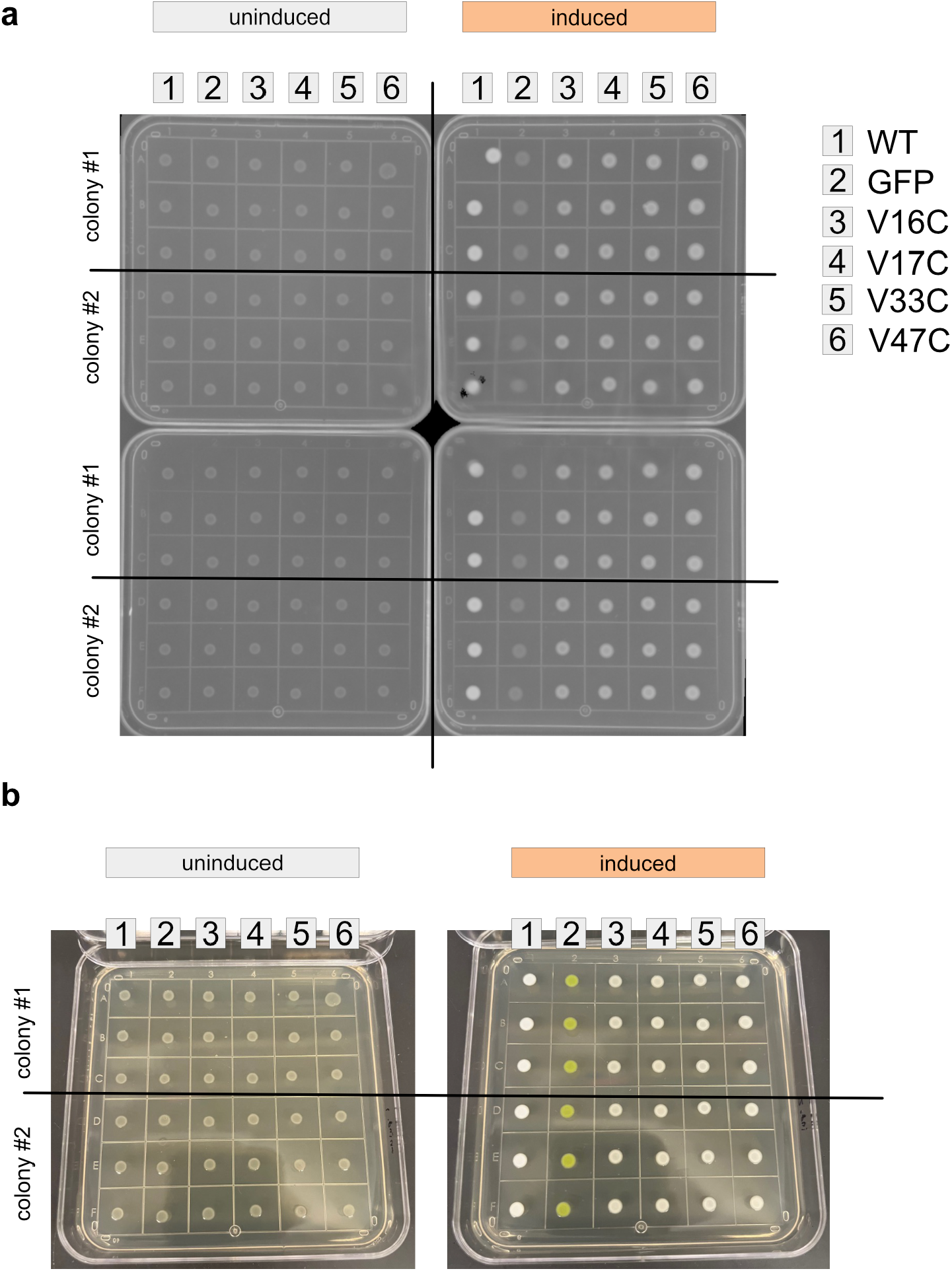
SonoCage candidate screen on patch plates. (a) Absorbance image of all four plates (two uninduced plates on left, two induced plates on right) taken 1 day after plating. On each plate, each column (labeled 1-6) denotes a different mutant or control. In each column, the first three rows are technical replicates made from the same original colony, and the next three rows are technical replicates made from another colony. (b) Phone camera image of two plates (one uninduced, one induced) taken 1 day after plating.

